# Accurate modeling of peptide-MHC structures with AlphaFold

**DOI:** 10.1101/2023.03.06.531396

**Authors:** Victor Mikhaylov, Arnold J. Levine

## Abstract

Major histocompatibility complex (MHC) proteins present peptides on the cell surface for T-cell surveillance. Reliable *in silico* prediction of which peptides would be presented and which T-cell receptors would recognize them is an important problem in structural immunology. Here, we introduce an AlphaFold-based pipeline for predicting the three-dimensional structures of peptide-MHC complexes for class I and class II MHC molecules. Our method demonstrates high accuracy, outperforming existing tools in class I modeling precision and class II peptide register prediction. We explore applications of this method towards improving peptide-MHC binding prediction.

## Introduction

T-cells play a crucial role in effecting and regulating the immune response in the context of infection, cancer, and autoimmunity. Their ability to recognize and respond to infected or abnormal cells is mediated by the interaction between T-cell receptors and peptides presented by MHC proteins on the cell surface. Predicting which peptides will bind to MHC, and understanding the properties of resulting peptide-MHC (pMHC) complexes such as immunogenicity and repertoires of cognate T-cell receptors, is essential for designing effective vaccines and immunotherapies. All of these properties ultimately depend on the structure of the peptide-MHC complex, making accurate *in silico* prediction of these structures an important task.

A number of pMHC structure prediction tools have been developed based on homology modeling, conformation sampling, and empirical energy minimization [1]. Protein modeling tools such as Rosetta [2] and MODELLER [3] were adapted to pMHC structure prediction in the context of immunogenicity prediction [4] and the modeling of pMHC-TCR complexes [5]. An example of an automated user-friendly tool is PANDORA [6], a MODELLER-based pipeline. Since the deep learning revolution in protein structure prediction [7,8], AlphaFold [8,9] (AF) has been applied to pMHC structure and binding prediction [10], and custom neural nets were built specifically for this task [11,12].

Our goal in this paper is to create and test an automated AlphaFold-based pipeline for pMHC modeling that would produce accurate models and be convenient to use. The main components of TFold, our pipeline, are paired pMHC template assignment, paired pMHC multiple-sequence alignments, and peptide register filtering with a sequence-based neural net. TFold works for peptides of different lengths in class I and class II structures with MHC alleles from human, mouse, and a few other species. It demonstrates high accuracy, for class I pMHCs significantly outperforming PANDORA [6]. For class II pMHCs, it outperforms state-of-the-art methods netMHCIIpan 3.2 and 4.0 [13,14] in peptide register prediction. Building on these results, we also explore applications of our pipeline to improved prediction of pMHC binding.

## Results

### A dataset of pMHC structures: peptide registers and geometric features

We searched the Protein Data Bank (PDB) [15] for entries that contain an MHC sequence, and collected structures representing 928 unique class I and class II peptide-MHC protein complexes. Most of these have human or mouse MHC alleles, but a number of structures with class I alleles for other species were also included (Table S1). In our workflow, the pMHCs for which at least one structure was deposited prior to the AlphaFold training cutoff date (2018-04-30) were assigned to a discovery dataset, which we used to explore pMHC features and optimize the pipeline’s hyperparameters. (No training of continuous parameters was done on the discovery dataset.) The rest of the data were assigned to a test set which was held out for validation. Many pMHC structures represent variants of commonly studied epitopes. To reduce redundancy, we clustered pMHCs by sequence distance (hierarchical clustering, tree cut at pMHC sequence mismatch equal to four) and chose one representative per cluster. The statistics of the resulting non-redundant dataset are plotted in Figure 1A. (Please see [Online Methods] for the details of data pre-processing.)

**Figure 1.**
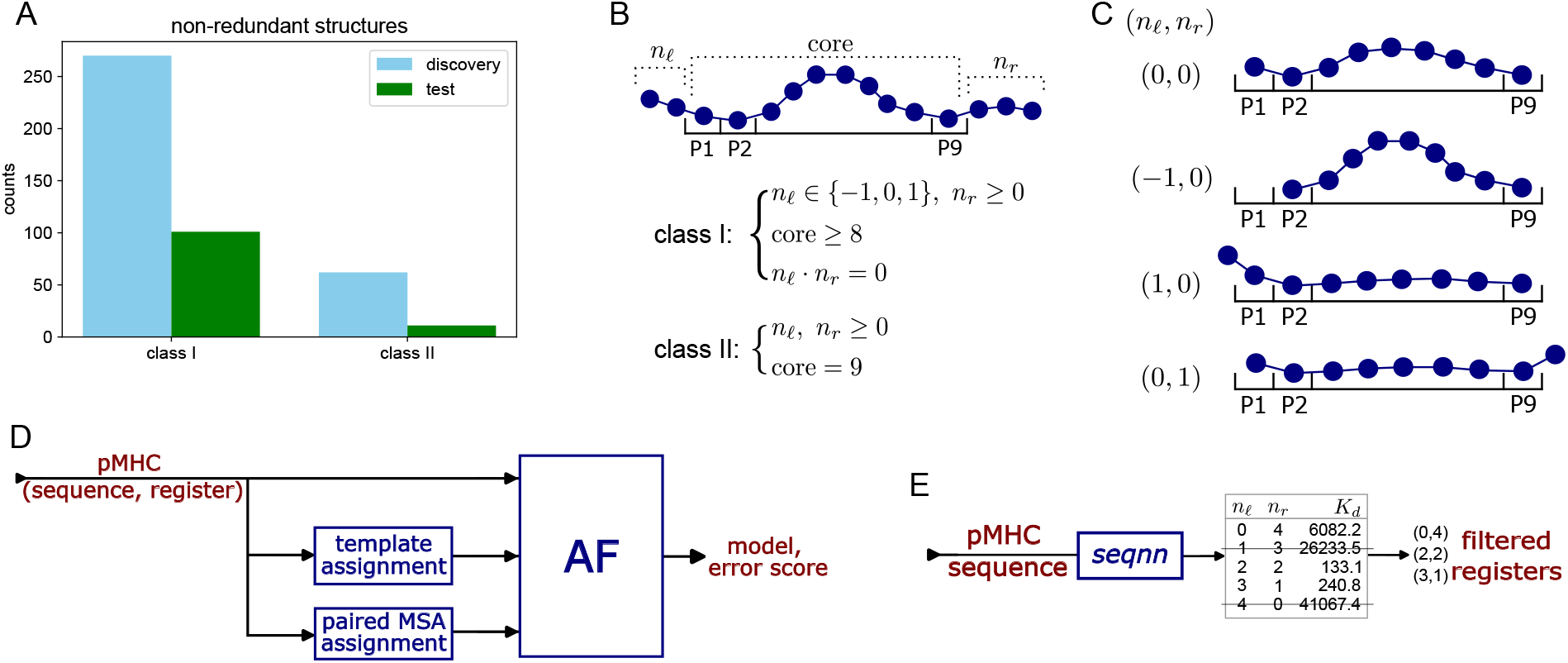
Structure dataset and the modeling pipeline. (A) Counts of non-redundant pMHCs in the discovery and test datasets, for class I and class II. (B) A schematic of a peptide position relative to the MHC binding groove. (For class I, positions P2 and P9 are the primary anchors. For class II, positions P1 and P9 are the two ends of the peptide core.) Peptide registers can be parameterized by the lengths of the C-terminal and N-terminal regions (*n_ℓ_*, *n_r_*). The sets of class I and class II registers observed in the discovery dataset can be characterized by a few simple rules. (C) The four registers that are possible for a class I 9-mer peptide, according to our register selection rules. (D) For a pMHC sequence and a choice of peptide register, our pipeline assigns templates and a paired peptide-MHC multiple sequence alignment. From these data, AlphaFold (AF) produces a model and an error score. (We use 100-pLDDT averaged over the peptide core as the score.) (E) A neural net *seqnn* predicts the pMHC dissociation constant *K_d_* for each peptide register. Only registers with *K_d_* within a certain factor of the lowest *K_d_* for a given pMHC are then considered in modeling.

For a pMHC structure, we define the peptide binding core (equivalently, peptide binding register) as the portion of the peptide that is positioned within the MHC binding groove. (In practice, we identify it by superimposing the structure onto a chosen template with a canonical 9-mer core, — PDB ID 3mre for class I and 4×5w for class II.) The binding register can be characterized by the lengths (*n_ℓ_*, *n_r_*) of the N-terminal and C-terminal peptide flanking regions (Figure 1B). Misidentifying the register in structure prediction leads to grossly incorrect models, and therefore, it is important to define the set of registers that are to be considered in modeling.

Among non-redundant class I pMHCs in the discovery dataset, around 95% of the structures assume the canonical register (*n_ℓ_, n_r_*)=(0,0) with the peptide ends tucked into the binging groove. The set of registers in the remaining 5% can be characterized by three simple rules (Figure 1B): (1) *n_ℓ_* ∈ { −1,0,1} and *n_r_* ≥ 0. (The case *n_ℓ_* = – 1 corresponds to the N-terminal residue of the peptide being the P2 primary anchor position); (2) the peptide core must be of length at least 8; (3) only one of *n_ℓ_* or *n_r_* can be non-zero. The third rule is likely probabilistic in origin: having a non-canonical terminal region is unlikely, and having both of them non-canonical is extremely unlikely. In fact, there is one structure in the discovery dataset that does not conform to these rules. It is a murine H2-Kd structure 5trz with register (−1,1), and it violates rule (3). In modeling class I structures, for peptide of any length we only consider registers that are allowed by the three rules above. Figure 1C illustrates the four registers that should be considered for a 9-mer peptide.

For class II structures that appear in the discovery dataset, the set of registers can be characterized by two rules (Figure 1B): (1) *n_ℓ_*, *n_r_* ≥ 0, and (2) the core length is nine. There exists one exception, a HLA-DRB1*01:01 structure 4gbx with register (−1,2), i.e., residue in position P1 missing. However, in modeling, we will assume that such exceptions are rare and will only consider registers allowed by the rules above. There exists *in silico* evidence [16] that some class II pMHCs adopt an unusual conformation, with the peptide overstretched or bulging out of the binding groove, and therefore core length different from nine. It is predicted that for some alleles, up to 10% of the peptides would adopt such conformation. We do not find any such examples in our discovery dataset, but it should be noted that it contains only four structures with alleles for which this behavior is predicted by [16] to be common.

A number of available class I and class II structures were not included in the discovery dataset because our register identification algorithm failed on them (Table S2). Most of these exceptions include peptide fragments, covalently modified peptides, or MHC bound to superantigens. Thus, we believe that our register selection rules of Figure 1B are fairly exhaustive. More details about the exceptions can be found in [Online Methods].

In modeling class I pMHC structures, our primary metric will be peptide RMSD (pRMSD), and more specifically, alpha-carbon peptide RMSD (*C_α_*-pRMSD). We define it by superimposing a model onto a true structure by MHC chains only, and then computing the error for the peptide. Unlike class I, for class II structures, the peptide core lies flat in the binding groove, while the terminal regions can adopt diverse conformations (Figures S3A,B). The pRMSD metric would then be dominated by errors in the flanking regions, which typically have few contacts with a T-cell receptor, except for residue P0 (Figure S3C,D). To focus on the most important geometric features, for class II models our primary metric will be peptide core RMSD (cRMSD), which only includes ten residues P0-P9 of the (extended) core of the peptide.

### TFold pipeline

The standard AlphaFold pipeline takes a list of protein sequences, builds a multiple-sequence alignment (MSA), and, optionally, finds templates for each chain in the PDB. The resulting data along with the chain sequences constitute the inputs to the AlphaFold neural net [8,9]. For modeling pMHC structures, this standard pipeline produces poor results [6,12]. Here we describe our custom AlphaFold-based pipeline TFold (Figure 1D,E) for modeling pMHC structures that is tailored to this particular problem and shows much better performance.

Poor results in using the standard AlphaFold pipeline for pMHCs can be attributed to the fact that no information about the relative position of peptide and MHC is provided to the neural net. In this situation, the task becomes an *ab initio* docking instead of a structure refinement. To remedy this, we provide custom templates that include both the peptide and the MHC chains, aligned to the whole pMHC sequence. The sequences are inputed as a single chain, and artificial 200-residue gaps between the chains are introduced into the AlphaFold internal residue indexing [17]. This way of sequence input allows the information about relative chain orientation in the templates to be used, unlike when inputing sequences as multiple chains in the multimer regime.

For template alignment, contact analysis, and other purposes, we introduce a peptide numbering system that assigns indices P1 and P9 to the N- and C-termini of the binding core. (If the register has *n_ℓ_*=-1 then residue P1 is missing and indexing starts with P2.) This numbering system is illustrated in Figure S1A. For class I structures with binding core longer than nine, we use single digit insertion codes after residue P5, and for shorter cores we omit residue P6, if needed. Index P0 with insertion codes is used for residues in the C-terminal flanking region.

Given a pMHC to be modeled, we consider all possible peptide registers subject to the selection rules described above and the pre-filtering process outlined in the next paragraph. A register choice defines a peptide numbering. In template assignment, this numbering is used to align the peptides, allowing when necessary to use templates with non-matching peptide length. MHC sequences are aligned according to the IMGT numbering [18]. Templates are then sorted by total pMHC sequence mismatch. AlphaFold takes four templates per run, and in default settings, we create five models per register, thus using the top 40 templates.

It is computationally costly to create models for all possible registers for a given peptide. Furthermore, for class I, we know that the canonical register is the right answer for ~95% of the structures, and we would like to impose this prior. One option is to use netMHCpan/IIpan [14] to predict the register, and only create models for this single register. However, as we assess below in this paper, these algorithms are not perfect at register prediction, especially for class II. Instead, we created our own sequence-based neural net *seqnn* for pMHC binding prediction. (It was trained on IEDB [19] data and netMHCpan/IIpan training data, separately for class I and class II. See [Online Methods] for details on architecture and training.) This algorithm gives access to *K_d_* predictions for each register separately. For modeling, we retain all registers that have predicted *K_d_* within a chosen threshold of the best predicted *K_d_* for a given pMHC. The threshold was set to 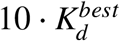 for class I and 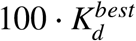 for class II. This cuts the average number of registers from 4.2 to 1.2 for class I, and from 6.9 to 4.2 for class II (data for the discovery dataset). On an NVIDIA A100 GPU, creating a single AF model takes about 30 seconds, and therefore, at five models per register, the average runtime is about 3 min and 10 min per class I and class II pMHC, respectively. For class II, the number of retained registers, and hence the runtime, may be higher for poor binders.

For a multiple-sequence alignment (MSA) to be useful, it needs to confer information about cross-correlations in peptide and MHC sequences. We construct paired pMHC MSAs, separately for class I and class II, from pMHCs identified as good binders in IEDB data and in netMHCpan/IIpan training sets [14]. MHC sequences are aligned according to their IMGT numbering [18], and peptides according to the numbering induced by their registers. For class I, peptide registers in the data were predicted using *seqnn*, and for class II, using TFold pipeline with no MSA. (See [Online Methods] for more details.)

Multiple models are produced for a single pMHC. To select the best one, and in particular, to predict the peptide register, we use the AlphaFold pLDDT score averaged over the peptide core. It is sometimes convenient to plot 100-pLDDT, to which we refer as the error score. We experimented with assigning different weights to pLDDT for anchor and non-anchor residues or adding scores that quantify template quality, but did not observe any improvement in model discrimination.

### Modeling class I pMHC structures

To evaluate the performance of the TFold pipeline for class I complexes, we model structures for non-redundant pMHCs from the discovery and test datasets. They mostly represent human loci HLA-A and B, as well as MHC alleles from mouse and other species, and a few HLA-C,E,G structures (Figure 2A). In modeling the discovery dataset, in order to ensure that the chosen templates are not closely related to the targets, we only allow templates from sequence clusters different from the target. For the test dataset, we allow arbitrary templates from the discovery dataset, which mimics the real life scenario where previously deposited structures are used to model a new target.

**Figure 2.**
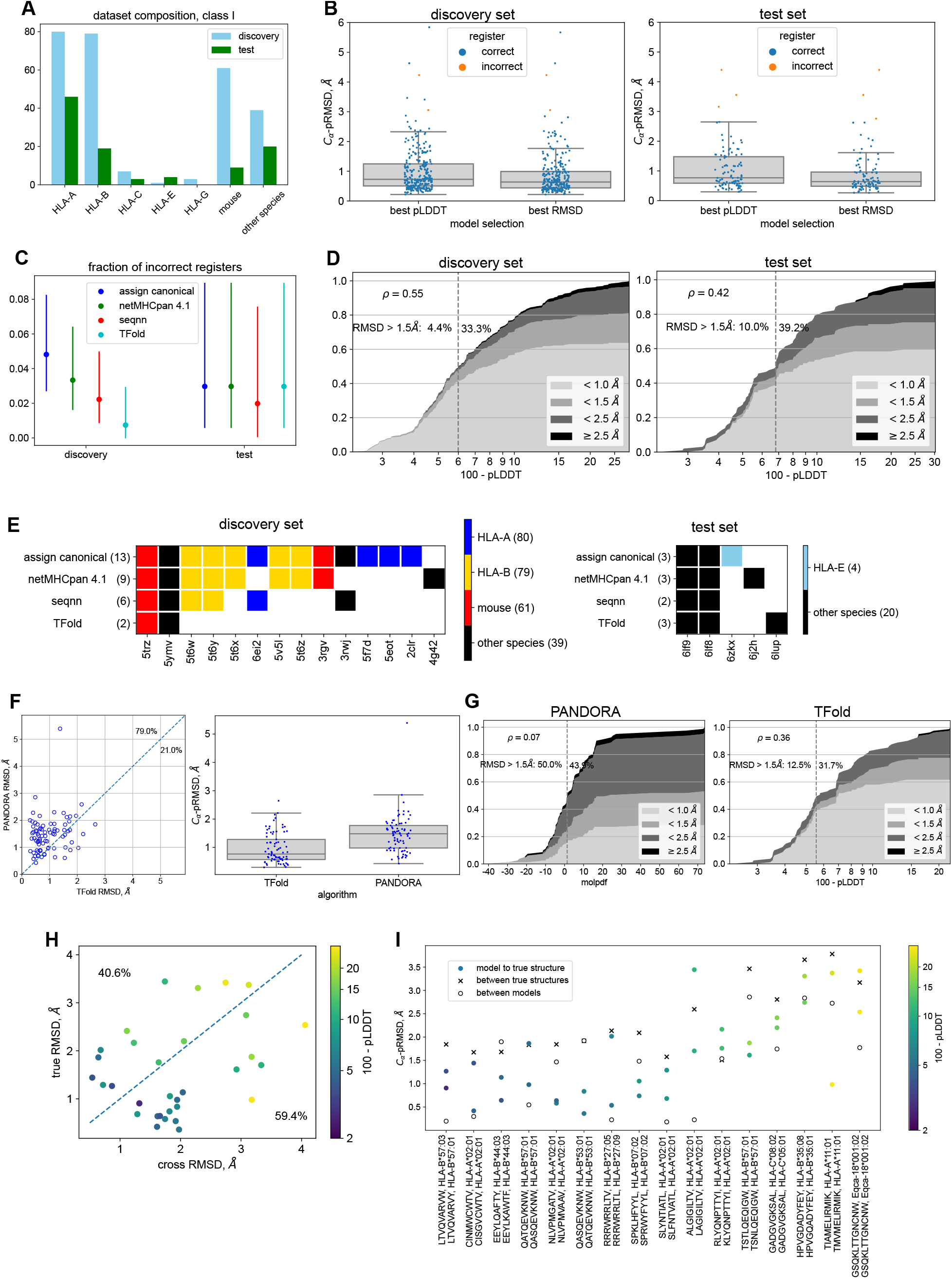
Results of pMHC modeling for class I structures. (A) Numbers of non-redundant pMHCs for different MHC loci and species in the class I dataset. (B) *C_α_*-pRMSD (alpha-carbon peptide RMSD) for TFold models in the class I discovery and test datasets. RMSD is computed upon superimposing MHC chains of the model and the experimental structure. Results shown for models selected by predicted LDDT (“best pLDDT”) and by peptide RMSD (“best RMSD”). Box plots show median value and first quartiles. (C) Fractions of incorrect predictions of peptide registers for class I models in the discovery and test datasets, for different methods. The first method (“assign canonical”) assigns the canonical register to all pMHCs. Bars show 95% confidence intervals (Agresti-Coull estimate). (D) Score vs accuracy plots for TFold models in the discovery and test datasets. Four accuracy groups of models based on *C_α_*-pRMSD are denoted by color: sub-angstrom (<1 Å), good (1-1.5 Å), poor (1.5-2.5 Å), and unacceptable (>2.5 Å). For every score cutoff (100-pLDDT plotted along the horizontal axis), the plot shows fractions of models in the four accuracy groups. A vertical dashed line marks the median score. For models below and above the median score, percentages of models with *C_α_*-pRMSD>1.5Å are shown to illustrate the score’s discrimination ability. Spearman’s *ρ* for the score vs RMSD is also printed on the plots. (E) Detailed diagram of register errors made by different algorithms on the discovery and test datasets. Rows correspond to algorithms, and columns to structures, with PDB IDs indicated below. Columns are colored by MHC locus and species. Each filled square indicates that the corresponding algorithm predicted the register incorrectly for the corresponding pMHC structure. (F) Comparison of *C_α_*-pRMSDs for class I models produced by PANDORA and TFold. Percentages in the left plot are fractions of pMHCs above and below the diagonal. Both algorithms were run on the subset of the test set that only includes human and mouse MHC proteins. (G) Score vs accuracy plots for models produced by PANDORA and TFold. (See caption for figure 2D for a description of such plots.) For PANDORA, the scores in the plot are values of the MODELLER [3] *molpdf* energy function. (H) TFold modeling results for the set of class I pMHC pairs that are similar in sequence but differ in geometry (“difficult pMHC pairs”). Each point in the scatterplot is a modeled pMHC, and the coordinates are *C_α_*-pRMSD of the model relative to the native structure (“true RMSD”) or to the experimental structure for the other pMHC in the pair (“cross RMSD”). Percentages indicate fractions of points above and below the diagonal. Points are colored by error score (100-pLDDT) of the models. (I) Details on the modeling results for the class I difficult pMHC pairs. Each column corresponds to a pair of pMHCs similar in sequence. (Some of them differ only by mutations in the MHC sequence, which are not shown.) Markers indicate *C_α_*-pRMSDs for models w.r.t. their experimental structures, between the two experimental structures, and between the models. Markers for model to native RMSDs are colored by the error scores, and average scores for each pair are used to sort the columns left to right. A perfect modeling algorithm would have low error for the models (colored markers near zero) and similar RMSDs between models and between true structures (crosses and empty circles overlapping).

TFold demonstrates sub-angstrom accuracy with median *C_α_*-pRMSD of 0.73 Å on the discovery dataset and 0.77 Å on the test dataset, showcasing its consistent performance (Figure 2B). (Median all-atom pRMSD is 1.55 Å and 1.77 Å, respectively.) If the best model is chosen for each pMHC instead of the model with the highest predicted accuracy, median *C_α_*-pRMSD improves slightly to 0.64 Å on both datasets.

On the discovery dataset, TFold predicts peptide registers better than netMHCpan 4.1, and also improves over *seqnn*, which is used for register pre-filtering (Figure 2C). TFold makes incorrect prediction for two structures, for which the other algorithms also fail (Figure 2E). One of these (5trz) has register (−1,1) and is the only exception in the discovery dataset to our register selection rules of Figure 1B. The other structure (5ymv) has a chicken MHC allele. We caution that the discovery dataset was used to choose hyperparameters for *seqnn* and TFold, and therefore these comparisons should not be over-interpreted. But notably, in all hyperparameter configurations that we have tested, TFold improved register prediction over *seqnn*.

On the test set, all algorithms show similar accuracy in register prediction (Figure 2C), however, these data include only three examples with non-canonical register. One of them (6zkx) has an HLA-E allele, with the register predicted correctly by all algorithms. The other two (6lf8, 6lf9) have swine MHC alleles, with registers predicted wrong by all algorithms (Figure 2E). TFold also predicts incorrectly a non-canonical register for a structure (6lup) with a shark MHC allele, while netMHCpan fails for a pMHC with a bat allele (6j2h).

Despite sub-angstrom median accuracy, TFold produces models of unacceptable quality for a substantial fraction of pMHCs. Many of these can be filtered out by setting a threshold on the predicted error score (100-pLDDT) which correlates well with *C_α_*-pRMSD (Spearman’s *ρ* equal to 0.55 on the discovery and 0.42 on the test data). We define four accuracy groups of models according to *C_α_*-pRMSD: sub-angstrom (<1Å), good (1-1.5Å), poor (1.5-2.5Å), and unacceptable (≥2.5Å). In Figure 2D, we plot fractions of pMHCs in these accuracy groups as a function of the score cutoff. This figure can guide the choice of a score cutoff in a modeling application according to the desired accuracy and fraction of pMHCs to be retained. For a low enough threshold, almost all modeled pMHC structures are sub-angstrom, while unacceptable models appear only at the highest values of the score. Separating the test set by median value of the score, only 10% of pMHCs in the top half are in the poor and unacceptable groups, compared to 39.2% in the bottom half, illustrating the score’s ability to enrich for high quality models. (For the discovery dataset, these fractions are even better, at 4.4% and 33.3%.)

We further explore what features of a pMHC are predictive of model accuracy (Figure S4A-D). Peptide length is the strongest predictor (Figure S4A), with median *C_α_*-pRMSD increasing from 0.46 Å for 8-mers to 2.14 Å for peptides longer than eleven residues. This is as expected, since longer peptides have a longer flexible loop region bulging from the MHC binding groove (Figure S3A). (Notably, this middle region that is hard to model is the most important one for TCR recognition, Figure S3C.) Next, we observe that structures with HLA-B alleles are on average harder to model than HLA-A (p=0.03, two-sided t-test) or HLA-C,E,G alleles (Figure S4B). Still, about half of HLA-B pMHCs are modeled with sub-angstrom accuracy. We do not observe any pattern in allele representation among HLA-B pMHCs with high vs low accuracy models. The lower accuracy for HLA-B could possibly be explained by worse template matching. Indeed, HLA-B templates on average have higher MHC sequence mismatch than HLA-A templates (Figure S4G). However, HLA-C templates have even higher mismatch, but modeling results for such pMHCs are no worse than for HLA-A. Furthermore, across all loci, MHC mismatch of the template is not a good predictor of modeling accuracy (Figure S4C). The last feature that we consider is peptide-MHC binding affinity. One may surmise that peptides that do not bind strongly have a less stable conformation and therefore are harder to model. A plot of netMHCpan-predicted binding affinity versus model accuracy demonstrates a weak relationship (Figure S4D). The set of peptide-MHC complexes that are selected for crystallization is enriched for good binders, and therefore a substantial fraction of such pMHCs for which netMHCpan predicts poor may be false negatives. It is therefore possible that the relation between binding affinity and modeling accuracy would be more pronounced if it were plotted for the experimentally-measured *K_d_*. Also, only a few pMHCs in our data are predicted to be poor binders.

### TFold significantly outperforms PANDORA, a MODELLER-based pipeline

We compare TFold to PANDORA [6], a state-of-the-art automatic pipeline for modeling pMHC structures. PANDORA pipeline is based on MODELLER [3] and comes with a structure database. Given a pMHC sequence, it finds a suitable template and performs anchor-restrained structure refinement. It uses netMHCpan to identify peptide anchor residues.

We compare the algorithms on the set of human and mouse structures from our class I test set. (PANDORA does not accept MHC alleles for species other than human and mouse.) For fair benchmarking, we removed from the PANDORA template database structures deposited after the AlphaFold training date. We remind that the same date cutoff was used to define our discovery dataset, which is the set of templates that TFold is allowed to use in this benchmark.

Peptide *C_α_*-RMSDs for models produced by the two algorithms are plotted in Figure 2F. TFold outperforms PANDORA on 79% of pMHCs (p-value < 10^−7^, Wilcoxon signed-rank test). TFold achieves *C_α_*-pRMSD of 0.77 Å, compared to 1.48 Å for PANDORA. (The original paper [6] reports median backbone pRMSD of 0.70 Å. This is likely due to the large number of close template-target matches, see Figure S3C of that reference.) Thus, our pipeline demonstrates a large and significant improvement in modeling accuracy.

PANDORA uses MODELLER’s score function molpdf to rank models. In Figure 2G, we compare its ability to discriminate structures to AlphaFold’s pLDDT. The Spearman’s correlation coefficient of molpdf with *C_α_*-pRMSD is 0.07, compared to 0.36 for the TFold score. Separating by the median molpdf score allows to filter out one out of three unacceptable models. For TFold, setting the threshold at median score filters out one out of one unacceptable models. Among PANDORA models with molpdf below the median value (highest predicted accuracy), 50% of the models are poor or unacceptable, which is more than 43.9% in the other half (lowest predicted accuracy). For TFold, these fractions are 12.5% and 31.7%, respectively. We conclude that TFold pipeline outperforms PANDORA in predicting model quality.

### Difficult pairs of class I pMHCs

Two pMHCs that are similar in sequence are typically close in structure. There exist exceptions to this rule — pMHCs where the change of one or a few residues substantially alters the peptide conformation. Such pMHCs may be of interest in the context of cancer. Indeed, if a single amino-acid mutant pMHC has peptide conformation significantly different from the corresponding wild type, it may be a good candidate for a neoantigen. Predicting effects of a minute sequence change is a challenging benchmark for a modeling algorithm.

In the class I discovery dataset, we identified sixteen pairs of pMHCs in which the two sequences belong to the same sequence cluster but the two structures have *C_α_*-pRMSD greater than 1.5 Å. (At most one pMHC in each pair was part of the non-redundant discovery dataset discussed in previous sections.) In some of these pairs, the pMHCs differ only in the peptide (1-3 substitutions), and in others, they have different but similar MHC alleles. We modeled this dataset withTFold, with the number of models per register increased from 5 (default) to 10 due to expected difficulty of the task.

Median *C_α_*-pRMSD on this dataset was 1.37 Å, which is quite a bit higher than 0.77 Å on the test dataset (Figure S4E). It is plausible that when a small sequence change can substantially disturb the structure, the conformation is not stable, making modeling harder. (Correcting for the different distribution of peptide lengths using per length RMSD from Figure S4A does not explain the difference.) Notably, the pLDDT score can still predict model quality well (Spearman’s *ρ*=0.70, Figure S4F). Let us define high predicted accuracy as error score below 6.8, the median score on the test set (Figure 2D). Restricting to models of high predicted accuracy, median *C_α_*-pRMSD becomes 0.84 Å, which is not far from 0.64 Å for high predicted accuracy models in the test set.

For a basic test of TFold’s ability to account for small changes in the sequence, for each model, we can compare its *C_α_*-pRMSD to the true structure (true RMSD) vs to the other structure in the pair (cross RMSD), see Figure 2H. For 59% of the models (19 of 32), true RMSD is smaller than cross RMSD, and if we restrict to models of high predicted accuracy, this number increases to 77% (10 of 13).

For a more detailed look into the modeling results, in Figure 2I, for each pMHC pair we plot *C_α_*-pRMSD between the true structures, between the models, and between the models and their respective true structures. Ignoring models with low predicted accuracy, two approximate patterns can be discerned in the data. For some pMHCs, the models are close to each other, i.e. TFold fails to account for the sequence difference. Both models can be close to one of the true structures, or they may interpolate somewhere in the middle. For other pMHCs, the models are far from each other and close to their respective true structures. These are modeling successes.

As an example, consider a pair of variant CMV pp65 epitopes NLVPMGATV and NLVPMVAAV, presented by HLA-A*02:01 (Figure S5A). For NLVPMVAAV, residue V6 is a secondary anchor, while the side chain of M5 is facing up and is available for TCR recognition. For the other variant NLVPMGATV, G6 has no side chain and instead M5 becomes a secondary anchor, turning away from the TCR. TFold correctly recognizes this conformational change, producing models that are far from each other and close to their corresponding native structures. Another example of a modeling success is a pair of HLA-B*53:01-presented HIV-1 Gag-Pol epitope variants QASQEVKNW and QATQEVKNW (Figure S5B). A single mutation adding a methyl group to the side chain in position three leads to a 1.92 Å change in the backbone and turns residue K7 towards the TCR. This is correctly reproduced by TFold models. However, for the same pair of peptides presented by HLA-B*57:01, the two TFold models generally follow the backbone of QASQEVKNW, failing to account for the sequence difference (Figure S5C). They both also have incorrect orientation of the K7 side chain. Similarly, for the pair of HCV NS3 epitope variants CINMWCWTV and CISGVCWTV presented by HLA-A*02:01, the two TFold models have similar backbones that are close to the native conformation for the second peptide, not being able to reproduce the difference in native structures. All models discussed in this paragraph have predicted error score below 6.8 (median score of the test set), and the score cannot distinguish the successes from the failures.

### Modeling class II pMHC structures

We modeled non-redundant class II structures from the discovery and test datasets. These structures represent all class II MHC loci for human and mouse, although notably almost half of the discovery dataset is HLA-DR, and the test dataset contains only eleven structures (Figure 3A).

**Figure 3.**
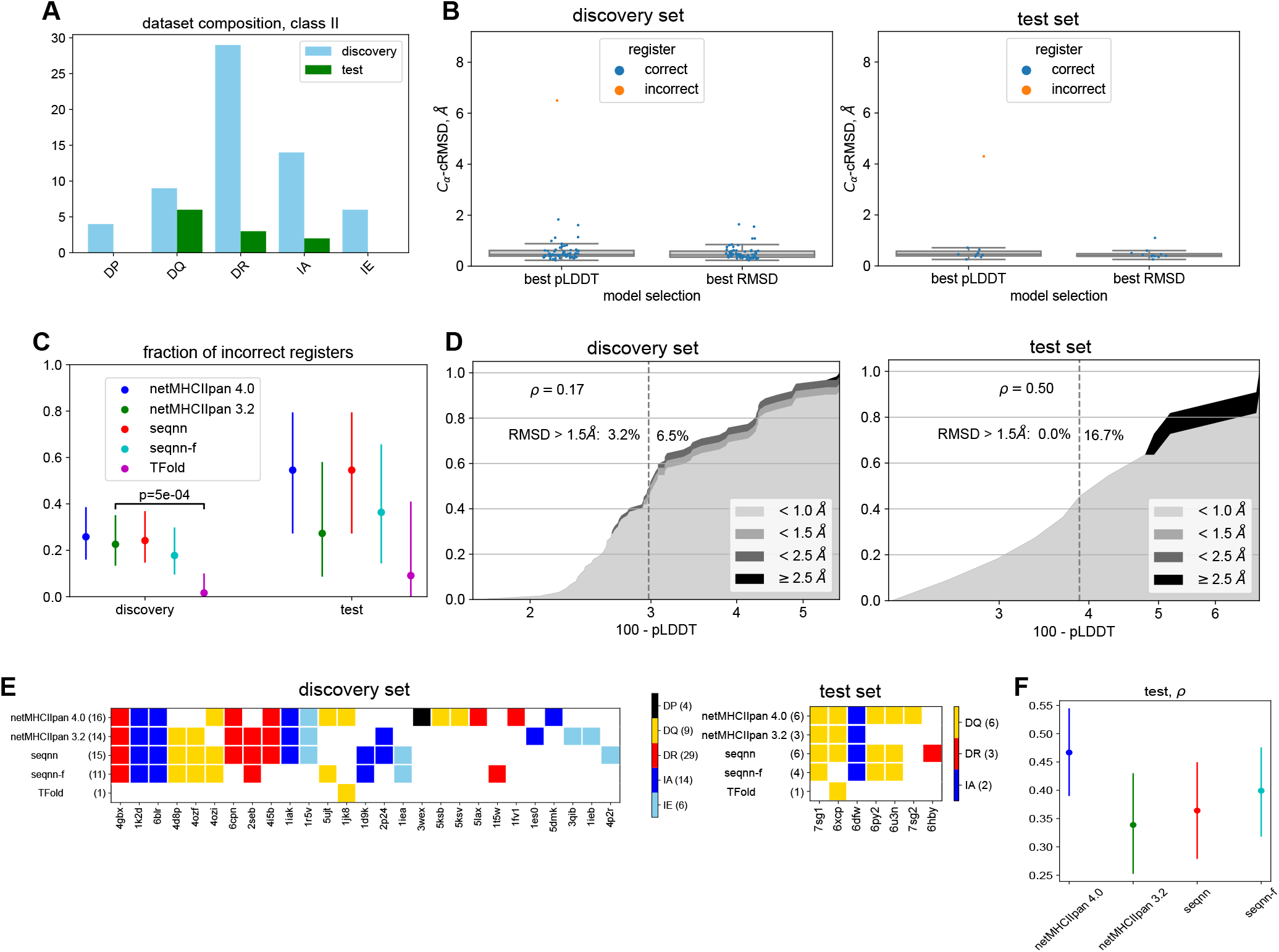
Results of pMHC modeling for class II structures. (A) Counts of non-redundant pMHCs for different MHC loci and species in the class II dataset. IA and IE denote the two class II mouse loci H2-IA and H2-IE. The dataset only includes human and mouse structures. (B) *C_α_*-cRMSD (alpha-carbon peptide core RMSD) for TFold models in the class II discovery and test datasets. RMSD is computed upon superimposing MHC chains of the model and the experimental structure, and only includes residues of the extended core P0-P9. Results shown for models selected by predicted LDDT (“best pLDDT”) and by peptide RMSD (“best RMSD”). Box plots show median value and first quartiles. (C) Fractions of incorrect predictions of peptide registers for class II models in the discovery and test datasets, for different methods. Bars show 95% confidence intervals (Agresti-Coull estimate). p-value is computed by Fisher’s exact test. (D) Score vs accuracy plots for TFold models in the discovery and test datasets. (See caption for figure 2D for a description of such plots.) (E) Detailed diagram of register errors made by different algorithms on the discovery and test datasets. (See caption for figure 2E for a description of such diagrams.) (F) Spearman’s *ρ* for the predicted vs measured pMHC dissociation constant (*K_d_*), for four different prediction algorithms. The test set here consists of *K_d_* measurements for 472 pMHCs with data deposited to IEDB after the netMHCIIpan 4.0 training date. Bars show the 95% confidence intervals estimated by bootstrap (1000 draws).

For the task of identifying the peptide binding register, we compared TFold to netMHCIIpan versions 3.2 and 4.0. (See Figure 3C. It also includes data for our pre-filtering neural net *seqnn* and its variant *seqnn-f*, which will be described below.) Of the two netMHCIIpan algorithms, version 3.2 performed the best, predicting incorrect register for 22.6% (14/62) of pMHCs on the discovery set and 27.3% (3/11) on the test set. (NetMHCIIpan errors are not due to a different definition of registers or shifted register patterns for different MHC alleles, see Figure S2D.) TFold makes a large improvement with 1.6% (1/62) and 9.1% (1/11) mistakes on the two datasets. For the discovery dataset, the difference with netMHCIIpan 3.2 is statistically significant (p-value 5e-4, Fisher’s exact test).

TFold register error rate on the test set is higher than on the discovery set. This could be just a fluctuation because the difference is not statistically significant (p-value 0.28, Fisher’s exact test). But there is another possible explanation. For all algorithms, register prediction for the HLA-DQ locus is harder (Figure S4H). The fraction of HLA-DQ pMHCs is higher in the test set than in the discovery set (Figure 3A), and this is enough to explain the difference: TFold error rates for DQ and non-DQ structures separately are consistent between the two datasets (Figure S4H).

There may be several reasons why structure prediction for HLA-DQ is harder. The HLA-DQ dimers are more diverse than HLA-DR due to the polymorphic *α*-chain. There are also less HLA-DQ than HLA-DR structures available for use as templates. In combination, these factors lead to worse template matching (Figure S4I). It has also been observed [20] in mass spectrometry data for naturally presented peptides that binding motifs for HLA-DQ alleles are hard to discern in the absence of the peptide exchange chaperon HLA-DM. If HLA-DQ proteins on their own have lower specificity, that would explain why identifying cores for HLA-DQ complexes is harder for a modeling algorithm that is not aware of HLA-DM. (This argument may also apply to the cores captured in experimentally determined structures, which we use as ground truth. Indeed, in crystallographic experiments peptides usually are not loaded via the native antigen presentation machinery.) We further note that TFold’s ability to predict class II binding cores is markedly reduced in the absence of paired peptide-MHC MSA input: the number of register errors increases from two to seven, six of which are in HLA-DQ examples. Thus, adding sequence information in the form of paired MSA is important for HLA-DQ register prediction.

The error score (100-pLDDT) is predictive of model accuracy (Figure 3D). The median score for the discovery dataset is 3.0, and the model with incorrect register appears at score 5.4 (ranked 61/62 by the score). In the test dataset, the model with register error appears at score 4.8 (ranked 8/11 by the score). However, only one out of nine HLA-DQ models in the discovery set and zero out of six HLA-DQ models in the test have score below 3.0, indicating that TFold has low confidence for HLA-DQ structures.

Alpha-carbon peptide core RMSD (*C_α_*-cRMSD) is plotted in Figure 3B. Median *C_α_*-cRMSD is 0.46 Å for both the discovery and the test set pMHCs. Median all-atom cRMSDs are 1.18 Å and 1.07 Å. Among pMHCs for which the register is predicted correctly, 93% in the discovery set and 100% in the test set are modeled with sub-angstrom accuracy (*C_α_*-cRMSD), highlighting the fact that flat peptide geometry of class II complexes (Figure S3B) is easy to model, once the register is identified.

In the description of the class II discovery dataset, we mentioned an HLA-DRB1*01:01 pMHC (PDB ID: 4gbx) that has peptide register (−1,2). This structure does not conform to the rules of Figure 1B, because one residue is missing at the N-terminus. Because of that, the register for 4gbx is predicted incorrectly by all sequence-based algorithms, but notably, TFold does not make a mistake (Figure 3E). The correct model is produced from the templates with register (0,1), demonstrating that AlphaFold is not confined to the immediate neighborhood of the template and may be able to recover a correct structure even when the template is misleading.

### Using TFold to identify registers at training improves performance of a sequence-based class II pMHC binding predictor

The data used in training sequence-based tools such as netMHCIIpan does not have labeled registers. During training, the neural net has to simultaneously solve the tasks of learning the motifs and selecting the registers where these motifs are the most prominent. Notably, the number of registers to consider is large, e.g. seven for a typical peptide length of fifteen. Fairly high register error (above 20% for both versions of netMHCIIpan and for *seqnn*, Figure 3C) attests to the difficulty of the problem.

We sought to leverage TFold advantage in class II register identification towards improving sequence-based binding predictors. We modeled top binders (*K_d_* up to 285nM) from the *seqnn* training set with TFold and selected pMHCs with models with high predicted accuracy (score below 1.3). This yielded a set of about 25000 pMHCs with predicted peptide register. We then trained a neural net *seqnn-f*, a version of *seqnn*, on the same data as before, but with the register fixed when a prediction is available. (The architecture and hyperparameters for *seqnn* and *seqnn-f* were optimized independently. The training process was two-step with additional register filtering. Please see Online Methods for the details.)

Seqnn-f outperformed seqnn but didn’t quite reach TFold accuracy in terms of register prediction, both on the discovery and test datasets (Figure 3C,E). To test its ability to predict the dissociation constant, we collected 472 pMHCs with measured *K_d_*, recently deposited in IEDB and not present in our or netMHCIIpan training set. On these data, seqnn-f demonstrated improvement over seqnn in terms of Spearman’s correlation between the predicted and measured *K_d_* values. Both seqnn and seqnn-f outperformed netMHCIIpan version 3.2 but not version 4.0 (Figure 3F).

Due to computational resource limitations, TFold modeling of pMHCs for the seqnn-f training set was performed without paired MSAs, which substantially decreases the accuracy of register prediction (register error rate 7/73 instead of 2/73 on the discovery plus test datasets). More importantly, netMHCIIpan version 4.0 leverages the massive dataset of naturally presented ligands [14], extending its training set by almost a factor of ten compared to version 3.2 (counting positive examples), while for seqnn and seqnn-f, we only used the binding constant assays. Competitive performance of seqnn-f relative to netMHCIIpan-3.2 suggests that with elution data included and with MSAs used in TFold runs, tools similar to seqnn-f may improve over state-of-the-art methods in class II pMHC binding prediction.

## Discussion

In this work, we developed TFold, an AlphaFold-based automated pipeline for modeling peptide-MHC structures. It can predict pMHC conformations for various class I and class II MHC loci from human, mouse, and a few other species for which templates are available (Table S1, Figures 2A, 3A). We analyzed peptide registers that occur in pMHC structures and incorporated this knowledge into the algorithm. The key elements of our pipeline are paired pMHC template assignment, paired multiple-sequence alignments derived from the binding data, and selecting subsets of peptide registers with a custom sequence-based neural net.

TFold shows competitive performance. For class I pMHC complexes, it achieves median peptide *C_α_*-RMSD of 0.77 Å, outperforming by a large margin a state-of-the-art MODELLER-based pipeline [6] (Figure 2B,F). We also demonstrated that the AlphaFold pLDDT score is a useful predictor of model quality (Figure 2D,G). We analyzed pairs of pMHCs that are similar in sequence but have divergent peptide geometry. Such pMHCs, when their peptides are derived from the human proteome and differ by a point mutation, may be interesting candidates for cancer neoantigens. Correctly translating a small change in sequence into an altered peptide conformation is a challenging task for a structure modeling algorithm. TFold is able to produce models that are closer to the true structure than to the structure of the other pMHC complex in the pair in 10 out of 13 cases, when restricted to pMHCs with high predicted accuracy. It is able to achieve some remarkable successes, such as catching the secondary anchor switch within the pair of very similar HLA-A*02:01-presented CMV pp65 epitope variants (Figure S5A). However, it also produces some fairly inaccurate models (Figure S5C), and the pLDDT score for them is not lower. There is room for improvement, and we suggest using this challenging benchmark in the evaluation of future pMHC structure modeling algorithms.

For class II pMHCs, TFold produces highly accurate models with median peptide core *C_α_*-RMSD of 0.46 Å (Figure 3B). (Our accuracy metric is focused on the peptide core residues P0-P9 because they make the most contacts with the MHC and T-cell receptors.) The main challenge in class II structure modeling is identifying the binding core, and in this task, TFold outperforms netMHCIIpan (Figure 3C). Moreover, setting a reasonable threshold for the predicted accuracy score allows to filter out register errors completely (Figure 3D). Predicting the binding core for peptides presented by MHC from the HLA-DQ locus is the most challenging both for TFold and for the sequence-based algorithms (Figure S4H). This can be related to worse template matching (Figure S4I) or to intrinsic properties of HLA-DQ proteins [20].

TFold ability to identify class II registers can be used to improve conventional sequence-based pMHC binding predictors by labeling registers in the training set. We demonstrated that this indeed improves our algorithm *seqnn* in terms of both register and dissociation constant prediction. (Figure 3C,F; the version of *seqnn* with TFold register filtering is labeled *seqnn-f*.) Notably, this way of leveraging the power of AlphaFold for binding prediction does not increase the runtime at inference. In training *seqnn* or *seqnn-f* in this paper, we did not use the elution data, which is critical for state-of-the-art prediction of binding [14], as we recapitulate in Figure 3F. Training a neural net with elution data included, and with registers in the training set labeled by TFold is a promising avenue for the improvement of class II pMHC binding predictors.

We envision that accurate *in silico* modeling of pMHC structures will facilitate the development of structure-based peptide:MHC and pMHC:TCR binding predictors. While this paper was in preparation, a related work [10] appeared that has substantial overlap with our work. It uses AlphaFold for accurate class I and class II pMHC structure prediction, making the crucial step of providing paired peptide-MHC templates. It does not use paired MSAs, which we observed to be helpful for accurate register identification in class II structures, but does AlphaFold finetuning on structural and binding data. It would be very interesting to see how AlphaFold finetuning would improve the accuracy of our pipeline. (For some direct evidence of such improvement in structure prediction see [21].) The focus of [10] is on binding prediction, while in the present paper, we focus on thoroughly evaluating the pMHC modeling abilities of AlphaFold, therefore we hope that our work will usefully complement these recent developments. In another exciting recent work [21], AlphaFold modeling is successfully applied to matching pMHCs to their cognate TCR repertoires, even for epitopes with no TCR training data.

Finally, we would like to highlight the difference between modeling class I and class II pMHC structures. In class I, apart from the rare cases (about 5%) of peptides with non-canonical binding registers, it is easy to position the primary anchor residues in the corresponding MHC pockets, and the challenge lies in modeling the peptide middle, including possible secondary anchors (Figure S3A). These middle residues are the most important for TCR recognition (Figure S3C), and even for 9-mer peptides, inaccurate models can substantially misrepresent the molecular features seen by a TCR (Figures S5B,C). For class II pMHCs, on the other hand, the peptide lies flat in the binding groove (Figure S3B) and is easy to model with consistent sub-angstrom accuracy (Figure 3B), once the binding register is identified and so long as we only focus on the peptide core. These observations may have implications for the accuracy of structure-based peptide:MHC and pMHC:TCR binding predictors.

## Acknowledgements

VM is grateful to Daniel Mattox, Damon May, Matthew Noakes, Ravi Pandya, and Jeremy Shaver for helpful discussions, and to Dario Marzella for help with setting up PANDORA. This work was supported by the NIH-NCI grant 5PO1CA087497-20.

## Data and code

TFold pipeline and the datasets used in this paper are available at https://github.com/v-mikhaylov/tfold-release.

## Online Methods

### Preparing sequence data

MHC protein sequences for species other than mouse were downloaded from the Immuno Polymorphism Database [22,23] on 2021-09-17. Mouse MHC sequences were manually curated from UniProt [24] and PDB [15]. Each sequence was aligned to an IMGT-numbered sequence [18] for the closest species and locus to establish IMGT numbering. For class I G*α*2 domains, the numbering was shifted by 1000 to distinguish from G*α*1 domains, e.g., a residue in the middle of an MHC alpha helix would be numbered 66 and 1066 for *α*1 and *α*2 helices, respectively. For consistency, we similarly shifted the IMGT numbering for class II G*β* domains. TCR *α*, *β, γ, δ* V- and J-gene sequences were downloaded from IMGT [25]. TCR V-regions were numbered according to the IMGT system [26].

### Preparing structures

Using the AlphaFold template search pipeline, we searched the PDB database (downloaded on 2022-07-05) for entries containing an MHC chain. In brief, MSAs were constructed for representatives of class I and both chains of class II MHC by applying Jackhmmer [27] on the Uniref90 [28] database. The MSAs were then used to search the PDB with hmmsearch [29].

For each chain fragment in a PDB file, protein type (MHC class and chain, TCR V and J regions, *β*2*m*), locus and allele were identified with a BLAST search. (Since allele assignments were done automatically, they may differ from alleles reported in PDB for the structures.) MHC and TCR segments were then realigned to numbered sequences using the BioPython [30] tool pairwise2.align to assign IMGT numbering. MHC chains were truncated to G-domains.

Class II MHC chains were matched together by proximity of *β*1-strand residues 4-11 and 1004-1011. TCR chains were matched together by proximity of fragments 49-53 and 115-118. TCRs and pMHC were matched into complexes by proximity of CDR3*β* to the peptide core. Chains were renamed into A (TCR *α* or *γ*), B (TCR *β* or *δ*), M (MHC *α*), N (MHC class II *β*), and P (peptide; see the next paragraph for peptide identification method).

Chain fragments that were not mapped to MHC, TCR or *β*2*m* were considered as potential MHC-bound peptides. For each MHC protein in a query structure, we superimposed the structure onto a reference structure (3mre for class I, 4×4w for class II) by MHC chains. For each residue in each candidate peptide fragment, we found the closest (C*α* distance) residue in the reference structure peptide. Fragments with minimal residue-pair distance below 2 Å were further considered. For them, we looked at consecutive residue pairs that mapped to residues (8,9) and either (1,2) or (2,3) in the reference structure peptide. If two such pairs were found, the corresponding fragment in the query structure was identified as a peptide bound to the MHC by which we superimposed. The same procedure allowed to identify peptide binding cores and therefore the registers.

This peptide identification procedure failed on 35 PDB structures that are listed in Table S2. We next comment on some of these exceptions. The reference structures to which the queries were superimposed have human MHC alleles, nevertheless, the procedure works surprisingly well even for MHCs from other species (Table S1), attesting to the evolutionary conservation of MHC geometry. However, Table S2 does contain one swine MHC structure. It also contains a human HLA-F structure, in which the peptide N-terminus is extended from the binding groove. It is known that HLA-F binds peptides differently than classic MHC class I proteins [31], and this geometry cannot be processed by our pipeline. Table S2 also contains an interesting example of a class II structure with the peptide bound in reverse. (There are two such structures, but in one of them the reverse orientation is enforced by a linker.)

In pMHC complexes made for crystallization, the peptide is sometimes connected to the MHC by a linker (1.3% of class I and 25% of class II PDB entries in our data). For such structures, we trimmed the corresponding peptide flanking region to zero in class I and to three residues beyond the binding core in class II complexes.

Contact counts for Figure S3 were computed as follows. Two atoms at distance *d* were considered in contact if *d* < 1.1 · (*r*_1_ + *r*_2_), where *r*_1_ and *r*_2_ are their van der Waals radii taken from [32]. For a pair of residues, the contact count is the number of pairs of heavy atoms in contact.

Structure manipulation, including superimposing and RMSD computations, was done using BioPython.PDB [33].

### Aggregating structure data and selecting pMHCs for benchmarks

The PDB often contains multiple structures for the same pMHC. Therefore, we merged structure records with identical pMHC sequences and peptide registers into pMHC records. (Before merging, gaps in peptide sequence were imputed from SEQRES entries, and gaps in MHC from the MHC sequence database.) For each pMHC record, the structure with the minimal number of missing peptide residues, with no linker (if available), and with the best resolution was chosen as a representative.

pMHC records were clustered into “sequence clusters” by sums of edit distances between peptide core sequences and between MHC sequences, separately for discovery and test datasets, with hierarchical clustering tree cut at distance 4. A representative pMHC was chosen for each cluster by the same criteria as in the structure per pMHC choice above. These representatives constitute our non-redundant discovery and test datasets of Figure 1A.

Separately, pMHC records were clustered by peptide backbone geometry (“geometry clusters”). For that, structures were transformed to the same frame by superimposing them onto the reference structures (3mre for class I, 4×4w for class II) by MHC chains. Vectors of peptide *C_α_* coordinates were collected and brought to the same dimension by restricting to residues P1-P9 (class I) or P0-P10 (class II) and imputing from neighboring residues. These vectors were clustered by k-means, with k set to the number of sequence clusters, and centroids initialized at average *C_α_* positions in the sequence clusters. In assigning templates in the TFold pipeline, in order to ensure template diversity, no more than one representative from each geometry cluster is allowed among templates for each pMHC-register pair.

For modeling and benchmarking, we used all non-redundant pMHCs from the discovery and test datasets, subject to the following restrictions: no missing residues in the peptide (class I) or in peptide positions P0-P9 (class II); no non-canonical residues in the peptide; no pMHCs with unstable peptide, i.e. for which available structures demonstrate more than one register.

The set of difficult class I pMHCs was chosen as follows. All records from the discovery dataset (not only non-redundant representatives) were filtered by the same rules as in the previous paragraph. Then we grouped pMHCs by sequence cluster and peptide length, and retained only groups with representatives of more than one geometry cluster. In each group, the matrix of *C_α_*-cRMSD distances was computed and the pair of pMHCs with the largest distance was retained. Further, we dropped pairs with peptide sequence mismatch greater than three or *C_α_*-cRMSD less than 1.5 Å. This resulted in the 16 pMHC pairs reported in the main text.

### Preparing peptide:MHC binding data

IEDB data for peptide:MHC affinity were downloaded on 2022-04-06. We only kept assays for peptidic antigens with no non-canonical amino-acids, presented by MHC alleles for which we have a sequence, with no mutations in MHC chains. Assay group “half maximal effective concentration (EC50)” was excluded. We further downloaded netMHCpan-4.1/IIpan-4.0 training data, which largely overlaps with the data from IEDB. For each unique pMHC, assays were merged by taking the geometric mean of the dissociation constants, and *K_d_* values were clipped to 1-50000 nM. Only pMHCs with peptides of length 8-15 (for class I) and 9-25 (for class II) were kept. IEDB pMHCs with the earliest assay deposited no earlier than 2020, and which do not appear in the netMHC training set, were set aside as a test set. The rest of the data, with the test set pMHCs excluded, were used as the training set. For training *seqnn* models, five training-validation splits of the training data were prepared.

### Paired multiple sequence alignments

Paired pMHC MSAs were constructed from the peptide:MHC affinity data in the training sets described above. MHCs were aligned in accordance with IMGT numbering, and peptides in accordance with their predicted registers.

For class I, registers were predicted using *seqnn*. We restricted to pMHCs with both measured and predicted dissociation constant below 100 nM, and further subsampled to no more than 100 peptides per MHC allele. This gave the final class I paired MSA of 8232 sequences.

For class II, register prediction with sequence-based neural nets is unreliable. Instead, we modeled the top 42413 binders (*K_d_* up to 285 nM) with TFold with no MSA. The top 10000 pMHCs by predicted accuracy were aligned for the MSA.

### Training seqnn and seqnn-f

Affinity-predicting neural nets *seqnn* were trained separately for class I and class II pMHC data. Their architecture is shown in Figures S1B, S2A. For each peptide register, one-hot encoded amino-acids are placed into input vectors according to the residue position number induced by the register choice. For class I, one residue beyond the binding core is included on each side. For class II, the 9mer binding core is provided to the network, as well as 3-neuron encodings of lengths of both flanking regions. The MHC is provided as a pseudo-sequence of 26 (class I) or 30 (class II) residues from positions with maximal peptide contact numbers in the discovery dataset. The input layer is followed by a number of fully-connected layers interspersed with batch normalization. The output neuron predicts logarithm of the dissociation constant. Minimal pooling over *K_d_* for different peptide registers is used to select a single value during training, but the network provides access to *K_d_* predictions for all registers on inference. Model architecture and hyperparameters were selected using the validation pMHC data and the discovery dataset of structures.

For class I, 40 models for each of the five train-validation splits were trained for 15 epochs. In the first 10 epochs, the canonical register was imposed for a randomly chosen half of 9mer peptides, to force a choice of the input neurons used for the binding core residues. We observed that the trained models clearly fall into two clusters with low/high register error on the discovery dataset, irrespectively of their *K_d_* prediction accuracy (Figure S1C). This indicates that some models learn a shifted pattern of anchors, for some or all MHC alleles. Therefore, we only retained models from the cluster with low register error by imposing a threshold of 23 errors on the discovery dataset. Geometric mean of the *K_d_* predictions of the resulting ensemble of 135 models is the output of *seqnn* for class I. The resulting predictor has slightly lower accuracy for *K_d_* (Figures S1D,E) and similar accuracy for register prediction (Figure 2C), compared to netMHCpan-4.1.

For class II, we first trained 40 models for 25 epochs for each of the five train-validation splits (200 models in total). Averaged register predictions of these models were used to label registers in the training set, and another 150 models were trained on these labeled data. Geometric average of predictions of these 150 models is the *seqnn* output. The resulting predictor performs similarly to netMHCIIpan-3.2 for *K_d_* prediction (Figures 3F, S2B) and similarly to netMHCIIpan-4.0 for register prediction (Figure 3C).

Our pipeline uses *seqnn* to pre-filter peptide registers before modeling. For each pMHC, we keep registers with predicted *K_d_* within a certain factor of the lowest predicted *K_d_* for that pMHC. If this factor is set too high, the filter would not eliminate any registers, and if it is set too low, true registers will be eliminated too often. This tradeoff is illustrated in Figures S1F and S2C for pMHCs from the class I and class II discovery datasets. We choose the thresholds at x10 and x100 for class I and class II predictors, respectively.

The network *seqnn-f* was trained on the same data as *seqnn*, but with registers partially labeled by TFold. We utilized models from the TFold run used to build the MSA (hence no MSA was used in the run), as described above. Therefore, models were available for 42413 pMHCs. The optimal neural network architecture was found to be similar to *seqnn*, but with more hidden neurons (four or five hidden layers with 512 neurons), and with dropout regularization (dropout fraction 0.6). Like for *seqnn*, the training procedure was two-step. In the first step, TFold models for pMHCs with predicted error score (100-pLDDT) below 1.3 were used to label registers on the training set, and 150 models were trained. In the second step, for pMHCs with no model of accuracy below 1.3, we labeled registers using the models from the first step (*K_d_* threshold set to x10). In this second step, 150 models were trained. The geometric mean of their output is the prediction of *seqnn-f*.

## Supplementary Figures and Tables

**Table S1.**
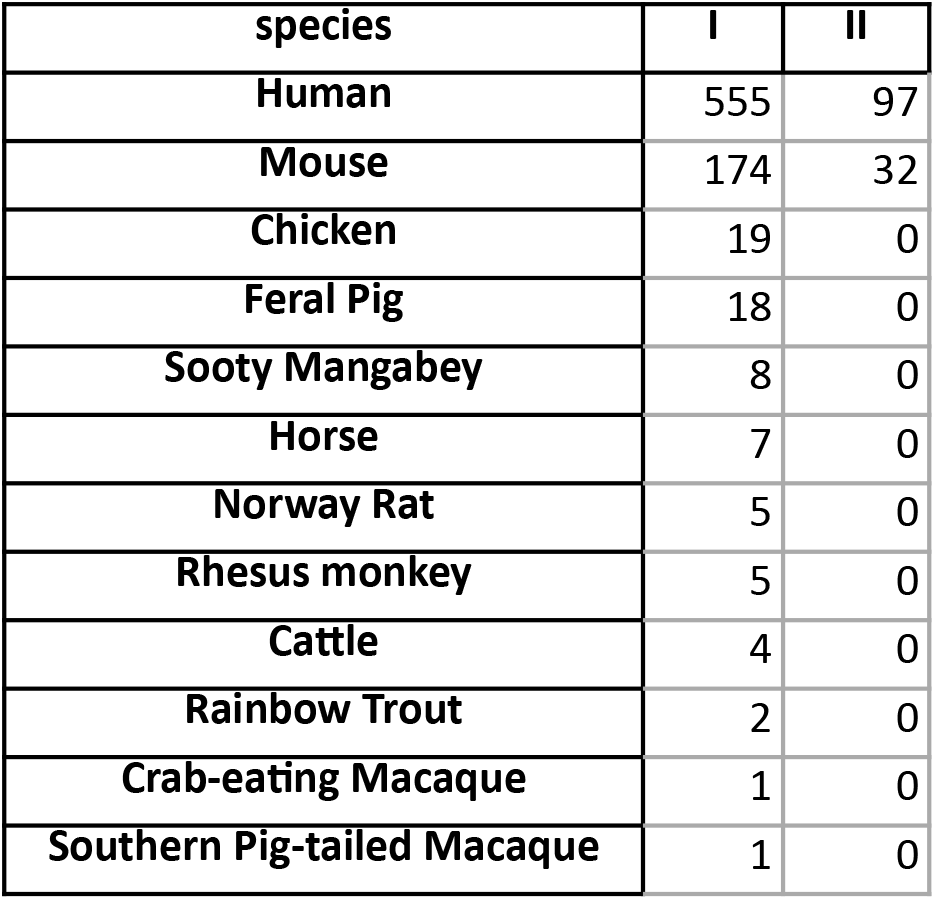
Unique pMHC counts in our structure database, by class and species. Species were assigned by aligning MHC sequences to the MHC sequence database and therefore may be approximate when the database does not contain the true species (e.g., trout instead of shark).

| species | I | II |
| --- | --- | --- |
| Human | 555 | 97 |
| Mouse | 174 | 32 |
| Chicken | 19 | 0 |
| Feral Pig | 18 | 0 |
| Sooty Mangabey | 8 | 0 |
| Horse | 7 | 0 |
| Norway Rat | 5 | 0 |
| Rhesus monkey | 5 | 0 |
| Cattle | 4 | 0 |
| Rainbow Trout | 2 | 0 |
| Crab-eating Macaque | 1 | 0 |
| Southern Pig-tailed Macaque | 1 | 0 |

**Table S2.** Structures for which our peptide register assignment script failed, with brief descriptions.

|  |  |
| --- | --- |
| **Class I** |  |
| Short peptides or peptide fragments | 1hsb, 2cii, 2x4n, 6vqy, 6vqz, |
| Missing peptide residues | 7kgt |
| Covalently modified peptides | 4uq2, 4zfz, 6iwg, 6iwh, 7byd, 7lfj, 7wj3, 7wt3, 6zky |
| Two peptide fragments (incl. structures with tapasin loops) | 5grg, 6jp3, 6gb5, 6gb7 |
| Peptide bound by disulphide bond | 5wes, 5wet, 6zkz |
| Peptide N-terminus conformation likely affected by linker | 4nhu |
| 8-mer with unbound C-terminus | 6vqd |
| HLA-F-bound peptide has N-terminus extending from the groove | 5iue |
| Unstable peptide conformation: three different registers within asymmetric unit | 4z78 |
| Swine MHC allele | 6a6h |
| **Class II** |  |
| Bound to superantigen | 1d5m, 1d5x, 1d5z, 1d6e, 1hqr |
| Bound to HLA-DM | 4fqx |
| Peptide bound in reverse | 3pgc, 4aen |

**Figure S1.**
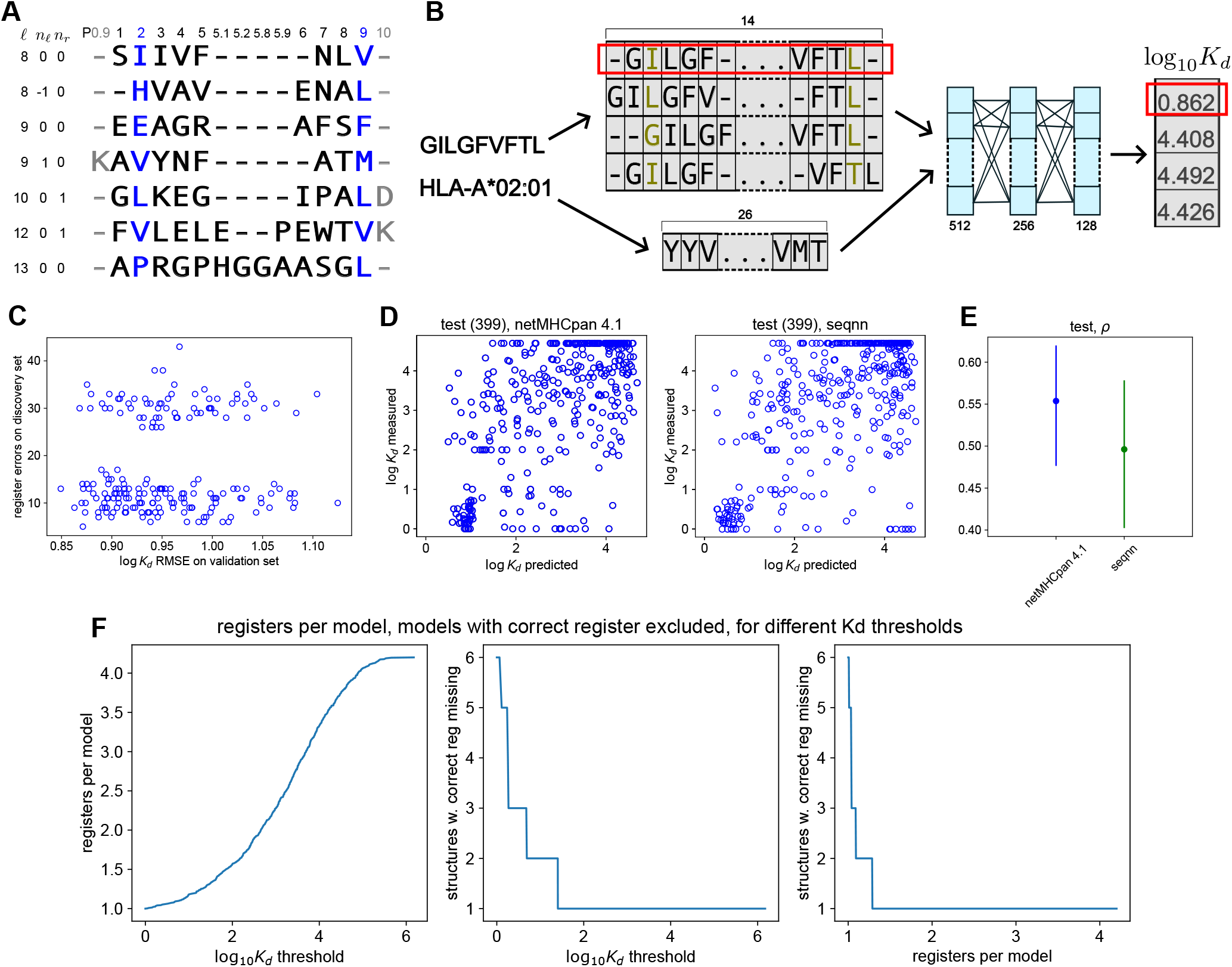
Peptide numbering and class I register pre-filtering. (A) A system for numbering peptide residues relative to the binding groove, illustrated for different peptide lengths and registers for class I pMHCs. Primary anchors P2 and P9 are colored blue, and flanking residues are colored grey. (For class II, the numbering system is the same except there are no positions with insertion codes after residue P5.) (B) Architecture of the class I *seqnn* neural network, showing peptide and MHC input. Each peptide register is processed independently. (C) Scatter plot showing error in log *K_d_* prediction vs error in register prediction for a set of independently trained class I *seqnn* models. RMSE for log *K_d_* is computed on a held-out validation set. The number of incorrect registers is for the class I discovery dataset. Two clusters of models with high/low register error are clearly visible. (D) Scatter plots for measured vs predicted log *K_d_* (*nM*) for netMHCpan and *seqnn*. The test set consists of *K_d_* measurements for 399 pMHCs with data deposited to IEDB after the netMHCpan 4.1 training date. (E) Spearman’s *ρ* for netMHCpan and *seqnn K_d_* predictions vs experimental values. (Same test set as in the previous panel.) (F) Varying *K_d_* threshold in register pre-filtering [Online Methods] changes the number of registers kept per pMHC and the number of pMHCs for which the correct register is erroneously filtered out. The three plots show the three quantities (threshold *K_d_*, register counts, structures with the correct register filtered out) plotted against each other for the class I discovery dataset.

**Figure S2.**
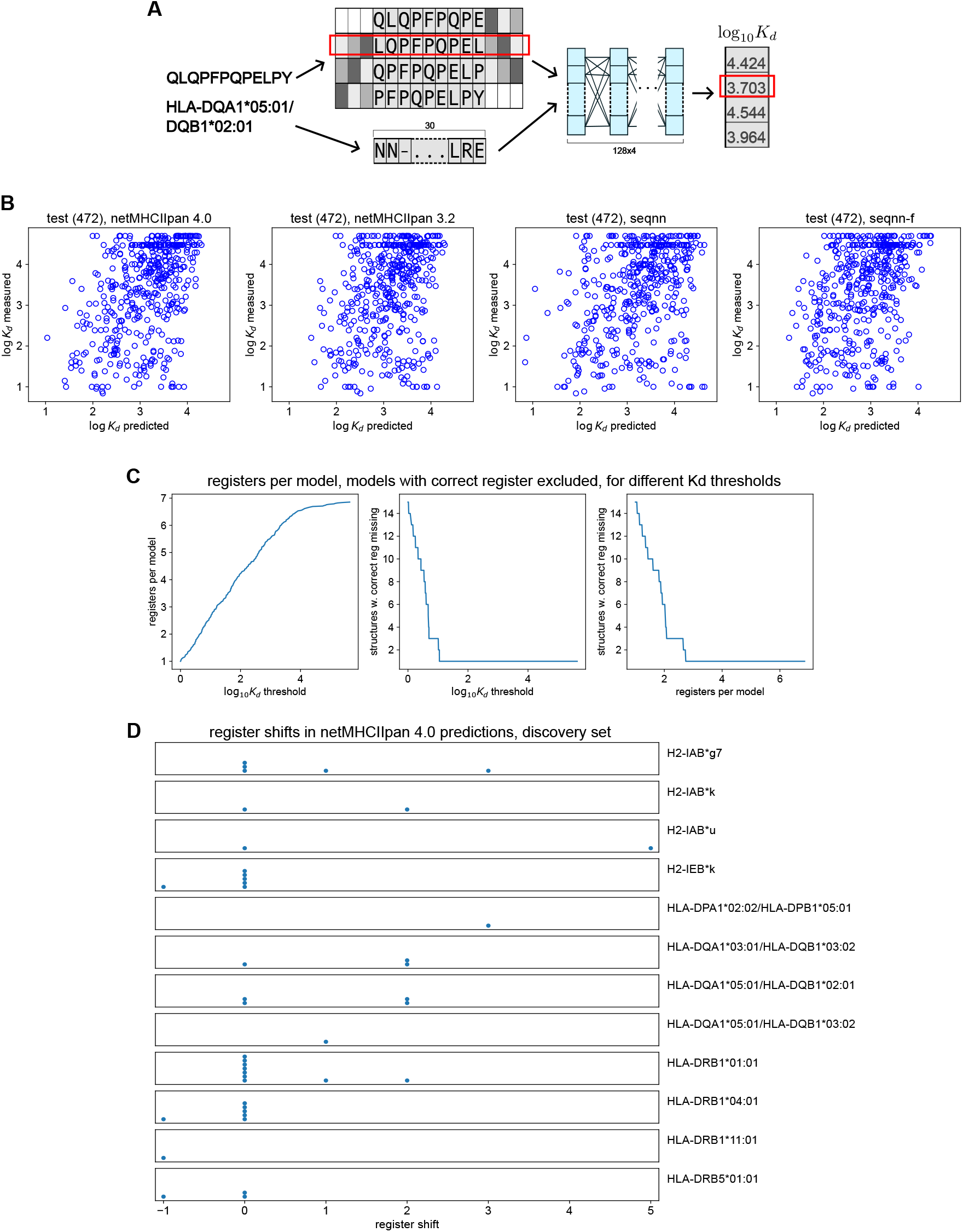
Class II register pre-filtering. (A) Architecture of the class II *seqnn* neural network, showing peptide and MHC input. Each peptide register is processed independently. Three bits on each side of the peptide encode length of the corresponding peptide terminus. (B) Scatter plots for measured vs predicted log *K_d_* (*nM*) for netMHCpan and *seqnn*. The test set consists of *K_d_* measurements for 472 pMHCs with data deposited to IEDB after the netMHCIIpan 4.0 training date. (C) Varying *K_d_* threshold in register pre-filtering [Online Methods] changes the number of registers kept per pMHC and the number of pMHCs for which the correct register is erroneously filtered out. The three plots show the three quantities (threshold *K_d_*, register counts, structures with the correct register filtered out) plotted against each other for the class II discovery dataset. (D) Shifts of netMHCIIpan 4.0-predicted peptide register vs the experimental register for structures in the class II discovery dataset. Each panel corresponds to an MHC allele, as shown on the right, and each datapoint is a pMHC structure. Only MHC alleles with at least one incorrectly predicted register are shown. For each such allele with more than one structure, at least two different shifts are observed.

**Figure S3.**
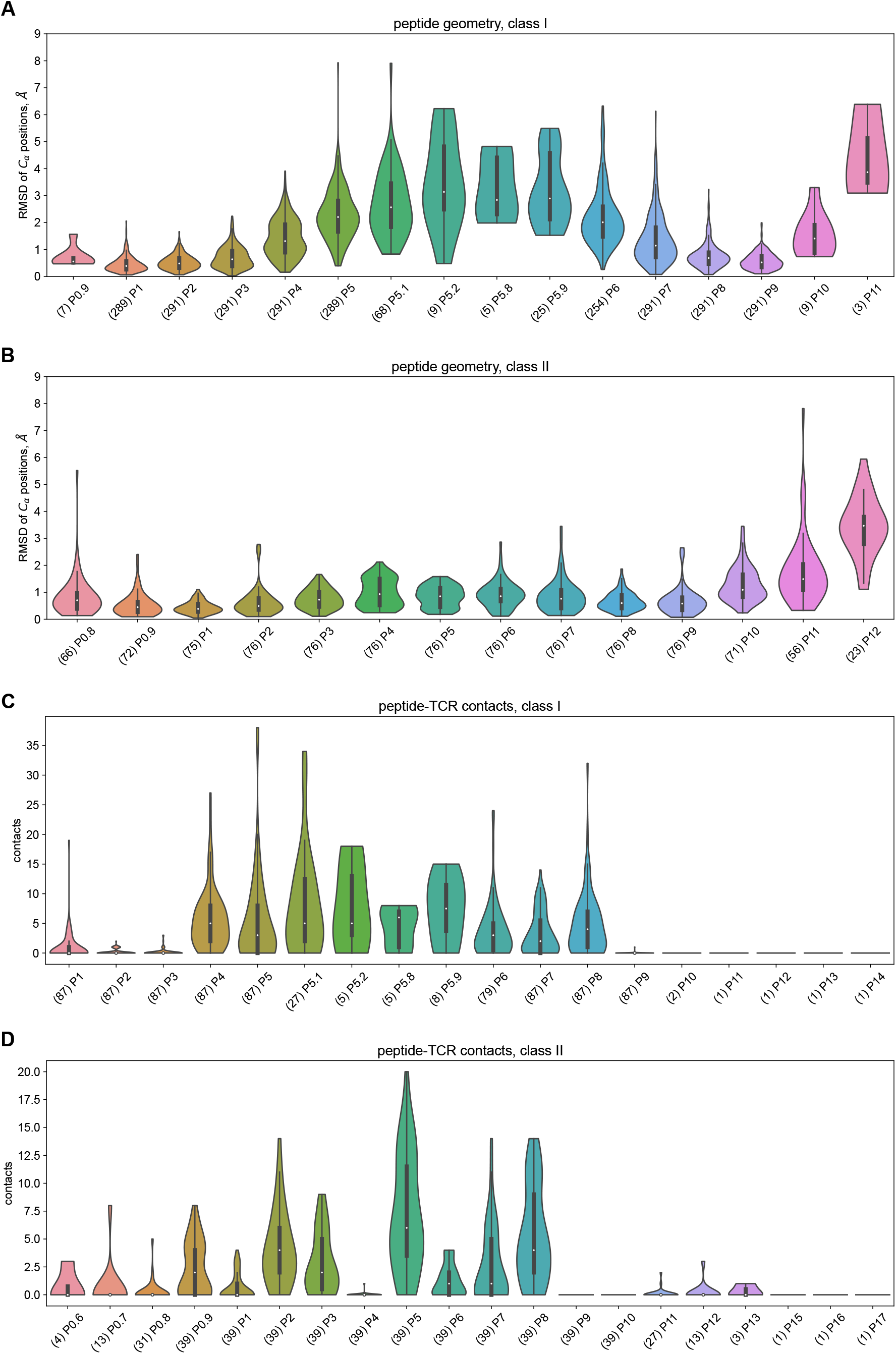
Peptide geometry and contact counts aggregated over the discovery dataset. (A) Distribution of *C_α_* positions for peptide residues, class I. Structures in the discovery dataset were superimposed by MHC chains, and for each peptide position, *C_α_* coordinate vectors were aggregated. The plot shows distributions of distances to the mean. Labels on the horizontal axis indicate peptide residue positions in our numbering system (Figure S1A) and (in brackets) the number of structures where a given position is present. (B) Distribution of *C_α_* positions for peptide residues, class II. (See caption for panel A.) (C) Distribution of TCR contact counts for different peptide positions, class I. Computed over structures in the discovery dataset that include a TCR. Labels on the horizontal axis are as in panel A. (D) Distribution of TCR contact counts for different peptide positions, class II. (See caption for panel C.)

**Figure S4.**
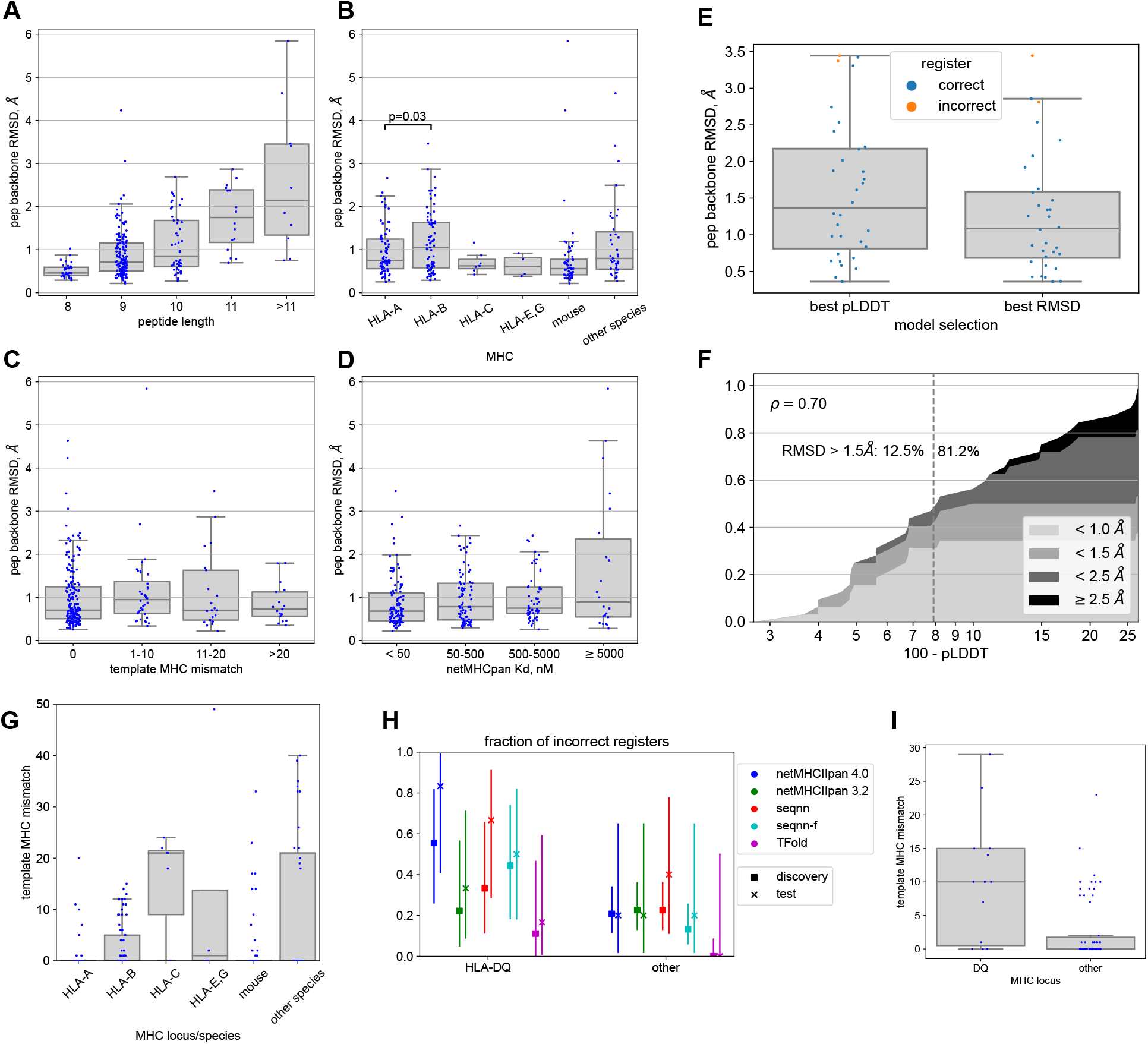
Additional details on structure modeling results. (A)-(D) Modeling accuracy as a function of different features, for the class I discovery dataset. The features include peptide length, MHC locus and species, MHC sequence mismatch of the best available template, and dissociation constant as predicted by netMHCpan 4.1. (E) Modeling accuracy for the class I difficult pMHC pairs (see also Figure 2H,I). The two box plots are for models selected by predicted accuracy (“best pLDDT”) and for the best models (“best RMSD”). (F) Score vs accuracy plots for TFold models for the class I difficult pMHC pairs. (See caption for figure 2D for a description of such plots.) (G) MHC sequence mismatch for the best template, for different MHC species and loci. Each point is a pMHC from the class I discovery dataset. (H) Fraction of incorrect registers as predicted by different algorithms for class II pMHCs from the discovery and test datasets, stratified by HLA-DQ vs all other loci. Bars show 95% confidence intervals (Agresti-Coull estimates). (I) MHC sequence mismatch for the best template, for HLA-DQ vs all other loci or species. Each point is a pMHC from the class II discovery dataset.

**Figure S5.**
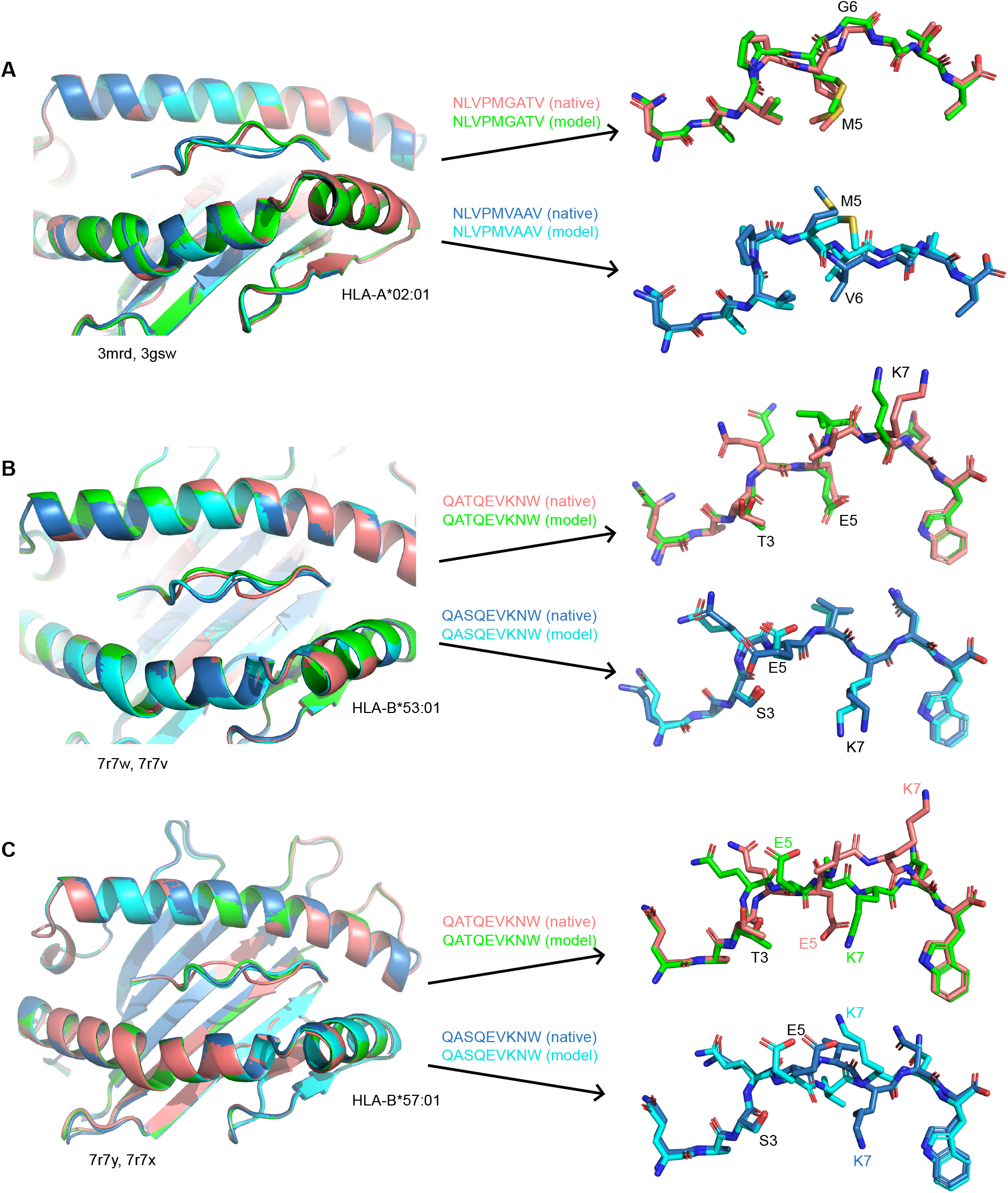
Examples of models superimposed on experimental structures, for three pMHC pairs from the set of difficult class I pMHC pairs. The structures are superimposed by their MHC chains. (A) HLA-A*02:01-presented CMV pp65 epitope variants NLVPMGATV (PDB ID: 3mrd) and NLVPMVAAV (PDB ID: 3gsw). *C_α_*-pRMSD between the two native structures is 1.84 Å, and model errors are 0.58 Å and 0.64 Å. (B) HLA-B*53:01-presented HIV-1 Gag-Pol epitope variants QATQEVKNW (PDB ID: 7r7w) and QASQEVKNW (PDB ID: 7r7v). *C_α_*-pRMSD between the two native structures is 1.92 Å, and model errors are 0.84 Å and 0.36 Å. (C) Same epitopes as in (B), but presented by HLA-B*57:01 (PDB ID: 7r7y, 7r7x). *C_α_*-pRMSD between the two native structures is 1.83 Å, and model errors are 1.86 Å and 0.98 Å.

